# User-friendly software for high-throughput processing of chlorophyll fluorescence images to assess plant stress responses: fvfmPy and fvfmR packages

**DOI:** 10.64898/2026.10.06.757212

**Authors:** Josef C. Garen, Pieter A. Arnold, Kristine Y. Crous

## Abstract

Chlorophyll fluorescence imaging is a powerful tool used to characterize abiotic and biotic stress response of photosynthesis in plants. However, extracting metrics such as *F_v_/F_m_* (maximum quantum yield of photosystem II) from chlorophyll fluorescence images remains a substantial bottleneck in large measurement campaigns, often requiring significant manual effort. To resolve this issue, we developed `fvfmPy`, a user-friendly program which enables rapid analysis of chlorophyll fluorescence images of arrays of leaf discs. `fvfmPy` uses algorithmic image segmentation to identify regions of interest and extract fluorescence metrics from tens or hundreds of leaf samples per image with little to no manual input. `fvfmPy` is developed in Python and includes a graphical user interface enabling intuitive parameter adjustment and real-time quality control checking. We additionally developed `fvfmR`, an R wrapper package for `fvfmPy`. *F_v_/F_m_* values extracted with `fvfmPy` were indistinguishable from those obtained using manual methods (*r*^2^ > 0.99) but required 94% less time to complete (13 s vs. 200 s per 100 measurements). `fvfmPy` and `fvfmR` were built to directly integrate with existing fluorescence software packages and can greatly increase throughput of fluorescence measurements to help address key challenges in agriculture and ecology.

**Highlight:** We developed a new, user-friendly tool to rapidly process chlorophyll fluorescence images. This software extracts maximum quantum yield and other metrics with minimal user intervention, enabling ultra-high-throughput image processing.

## Introduction

Developing new climate-resilient crops and understanding how global change affects ecosystem function are key research goals that necessitate understanding plant stress response at scale (Doughty *et al*., 2023; Eckardt *et al*., 2023; Posch *et al*., 2026). Consequently, there is growing demand for high-throughput phenotyping approaches to rapidly characterize stress responses across large numbers of species, accessions, or genotypes (Furbank and Tester, 2011; Fiorani *et al*., 2012; Granier and Vile, 2014; Yang *et al*., 2020). While screening efforts carried out across genetically diverse germplasm or novel crop species can aid in yield improvement and selection for tolerance to multiple stressors, such efforts have high data processing demands (Araus and Cairns, 2014). Similarly, characterizing the vulnerability of ecological communities to climate warming requires large-scale collection and processing of physiological data representing many species and multiple scales of variation (Geange *et al*., 2021; Wilkening *et al*., 2024; Cavender-Bares *et al*., 2025). The rapidly increasing volume and complexity of data from novel digital phenotyping approaches necessitate automated, reproducible, and scalable workflows that ensure consistency, transparency, and efficiency in data analyses (Humplík *et al*., 2015; Yang *et al*., 2020).

Chlorophyll fluorescence is one such modern phenotyping tool that is widely employed to quantify photosynthetic performance and plant stress responses in both crops and ecological communities (Baker and Rosenqvist, 2004; Humplík *et al*., 2015; Pérez-Bueno *et al*., 2019). Chlorophyll fluorescence has a wide variety of applications, including measurement of heat tolerance and thermal load sensitivity of photosystem II (Arnold *et al*., 2021, 2026, Preprint; Geange *et al*., 2021). Additionally, fluorescence is commonly used as a screening tool in agronomic applications to assess tolerance in crops to a wide variety of abiotic stressors including drought, cold, and salinity (Sayed, 2003; Baker and Rosenqvist, 2004; Arief *et al*., 2023). Chlorophyll fluorescence imaging, in particular, offers great potential for sensitive, high-throughput screening of many individuals simultaneously and offers spatially explicit information on tissue-level fluorescence (Kautsky and Hirsch, 1931; Schreiber *et al*., 1986; Murchie and Lawson, 2013). Image-based phenotyping has become an essential component of plant research, enabling non-destructive measurements of plant morphology and physiology across a large number of species (Baker and Rosenqvist, 2004; Humplík *et al*., 2015). However, image processing and analysis often requires substantial manual intervention including decisions on image segmentation, quality control, and trait extraction (Minervini *et al*., 2015; Tsaftaris *et al*., 2016). Thus, analysis of chlorophyll fluorescence images represents a major processing bottleneck in phenotyping pipelines, restricting its potential as a high-throughput methodology.

The development of automated image-analysis pipelines can provide substantial time savings, as well as improving analytical consistency and reproducibility between experiments and users (Tsaftaris *et al*., 2016; Pottinger *et al*., 2025). General-purpose image analysis programs such as ImageJ and PlantCV offer powerful and flexible toolkits, but using these programs to develop and implement automated pipelines for processing large volumes of data remains challenging for users (Katabuchi, 2015; Schuhl *et al*., 2026). This is especially true when analysing data packaged in proprietary or otherwise complex file formats, as is often the case with imaging fluorometry systems (Schneider *et al*., 2019). Purpose-built applications can help resolve these issues by tailoring image processing algorithms and file handling to specific use-cases and removing unneeded software complexity and computational overhead. For example, purpose-built automated image analysis programs have been developed for the detection of leaf area and shape (Bylesjö *et al*., 2008; Katabuchi, 2015), leaf venation traits (Xu *et al*., 2021), rosette fluorescence in *Arabidopsis* (Schneider *et al*., 2019), whole-plant traits in maize (Souza and Yang, 2021), and many other applications, enabling rapid and reproducible data analysis in specific domains. Still lacking, however, is a purpose-built application for analysing chlorophyll fluorescence images of leaf disc arrays, an increasingly common approach for high-throughput measurement of chlorophyll fluorescence (Arnold *et al*., 2021; Bison and Michaletz, 2024). Developing a purpose-built tool for this application is especially timely as chlorophyll fluorescence imaging is rapidly being integrated into large-scale phenotyping platforms and efforts to understand stress responses of plant species in a future climate (Harbinson *et al*., 2012; Rousseau *et al*., 2015; Yang *et al*., 2020).

Here, we present `fvfmPy` and `fvfmR`, user-friendly pipelines for rapid analysis of chlorophyll fluorescence images. This new software can automatically segment chlorophyll fluorescence images, detect regions of interest, and extract fluorescence metrics with little manual intervention, drastically reducing the time required to analyse fluorescence images of leaf arrays.

The software was developed in Python with an associated R wrapper package and is freely available for use and modification. In addition, `fvfmPy` and `fvfmR` easily integrate with existing packages for analysis of chlorophyll fluorescence data, enabling ultra-high-throughput data processing and analysis from measurement to final product.

## Materials and Methods

### Plant material

We grew 28 species of *Eucalyptus* and *Corymbia* saplings at Western Sydney University, Hawkesbury Campus (33°37’14“S, 150°45’14”E) during summer and autumn 2025. Plants were grown in 5 L pots in a mixture of 50% perlite, 30% sand, and 20% peat, and kept well-watered in a shadehouse when not being measured. Six to seven individual plants per species plants were transported to the laboratory on the day of measurement. Measurements took place on 2-5 June 2025 resulting in 360 arrays of 28 to 47 leaf disks per array. A subset of 40 chlorophyll fluorescence images including all species and comprising 1799 total measurements were used in the validation test (see *Validation dataset* below). Data from a subset of five de-identified species (SP1-5) were used in the thermal load sensitivity example (see *Example application: Thermal Load Sensitivity of photosystem II* below).

### Fluorescence measurements

We measured chlorophyll fluorescence, photosystem II heat tolerance, and thermal load sensitivity in our target species following the approach described in (Arnold *et al*., 2026, Preprint). In brief, small (c. 0.7 cm^2^) leaf discs were excised from healthy, fully expanded leaves and affixed in rectangular arrays to cardstock. Leaf arrays were dark adapted for a minimum of 30 minutes, then chlorophyll fluorescence measurements were acquired using a pulse-amplitude modulation (PAM) imaging fluorometer (Max-Imaging-PAM; Heinz Walz GmbH, Effeltrich, Germany). This measurement is referred to as the “initial” measurement. Leaf arrays were then sealed into clear plastic envelopes and placed into temperature-controlled water baths. Twenty-eight total heat dose treatments were applied, comprising seven temperature treatments (30, 36, 40, 44, 48, 52, and 56 °C) and five duration treatments (5, 15, 30, 60, or 120 minutes). Not all possible combinations of temperature and duration were employed. As heat stress naturally occurs in the light, an LED light source was supplied above the water baths with a photosynthetic photon flux density (PPFD) of 1100 μmol m^-2^ s^-1^ at the surface of the water bath. The light level that reached leaves in the water bath was 800-850 μmol m^-2^ s^-1^. Following treatment, leaves recovered in saturating light of 750-800 μmol m^-2^ s^-1^ at room temperature for 90 minutes. Leaves were then dark adapted again for 30 minutes and chlorophyll fluorescence was re-measured; this is the “post” measurement. Finally, leaves were placed in darkness overnight for 24 hours, and fluorescence was measured a third time; this is the “overnight” measurement.

### Overview of fvfmPy and fvfmR software packages

We developed the software packages `fvfmPy` and `fvfmR` to expedite image processing of chlorophyll fluorescence measurements. The core program, `fvfmPy`, is written in Python and is functional as a standalone software product. This software can also be run from within a native R programming environment as we also developed `fvfmR`, a wrapper package written using the `reticulate` R package. The overall workflow of `fvfmPy` is illustrated in Fig. 1. The workflow begins by collecting fluorometry images using rectangular arrays of leaf discs. Once the fluorometry images are collected, the user can process the images using `fvfmPy.` The graphical user interface of `fvfmPy` is illustrated in Fig. 2. Within the software, the user begins by specifying a directory containing fluorescence files in .pim (proprietary Walz format) or multi-frame .tif format. The software loads each fluorescence file sequentially and then attempts to automatically identify leaf disc regions of interest (ROIs) and extract fluorescence metrics (*F_o_, F_m_,* and *F_v_/F_m_*). Optionally, the software also allows the user to rotate or crop the fluorescence image if desired, or to adjust image segmentation parameters, as this may improve automated ROI identification in some cases. The user is presented with the automatically identified ROIs within the graphical user interface to allow rapid quality checking and manual adjustment if needed. Finally, the extracted fluorescence metrics are collated and output in a long-format .csv file for further processing steps (Table 1). Details of the automatic image segmentation algorithm and optional manual processing steps are given in the next two subsections.

**Figure 1:**
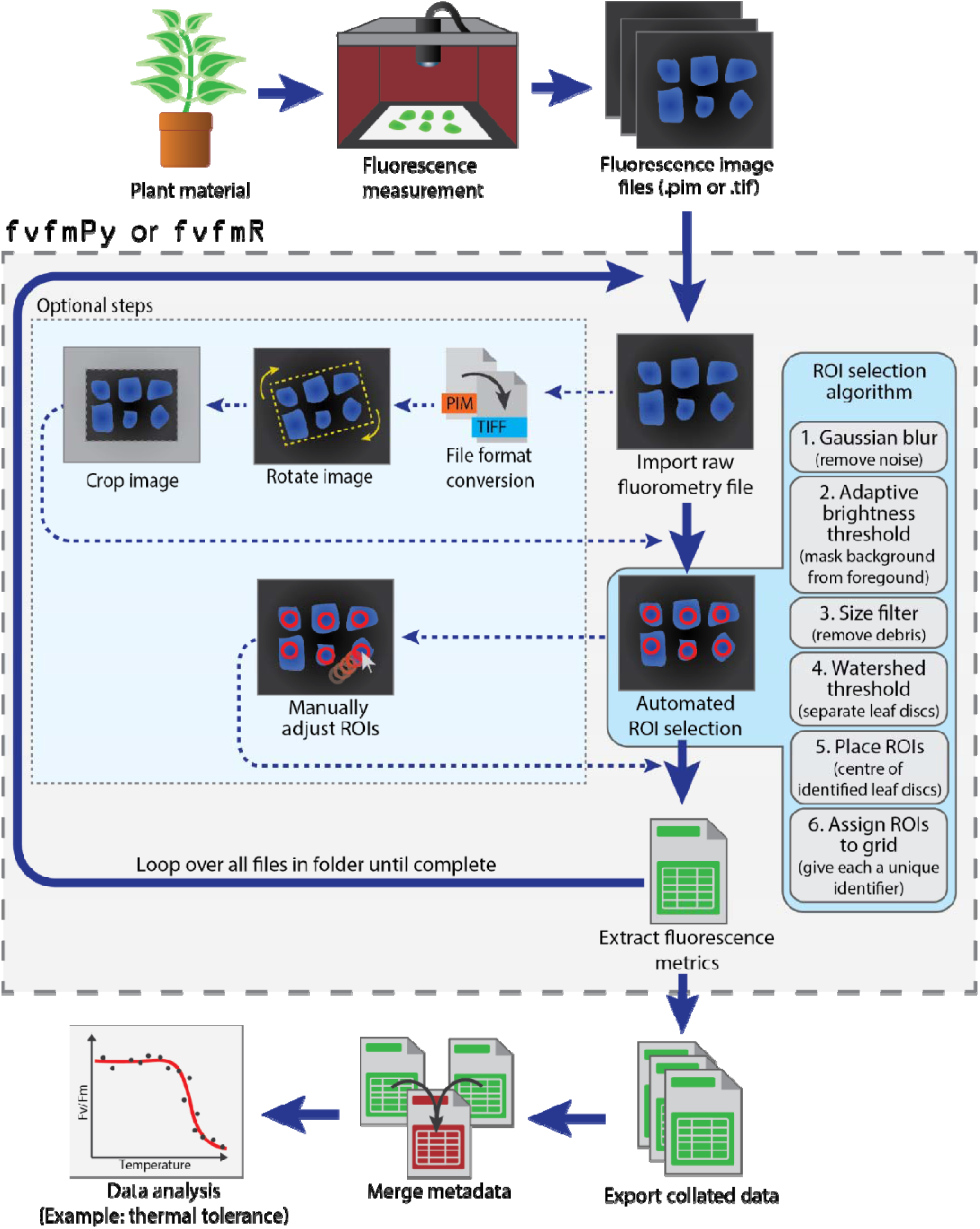
Workflow diagram for fvfmPy or fvfmR. This figure illustrates the use of `fvfmPy` or `fvfmR` in a large chlorophyll fluorescence measurement campaign. Internal data processes steps in `fvfmPy` are highlighted in the dashed box. `fvfmPy` and `fvfmR` enable rapid processing of chlorophyll fluorescence images, a common bottleneck in high-throughput data collection campaigns.

**Figure 2:**
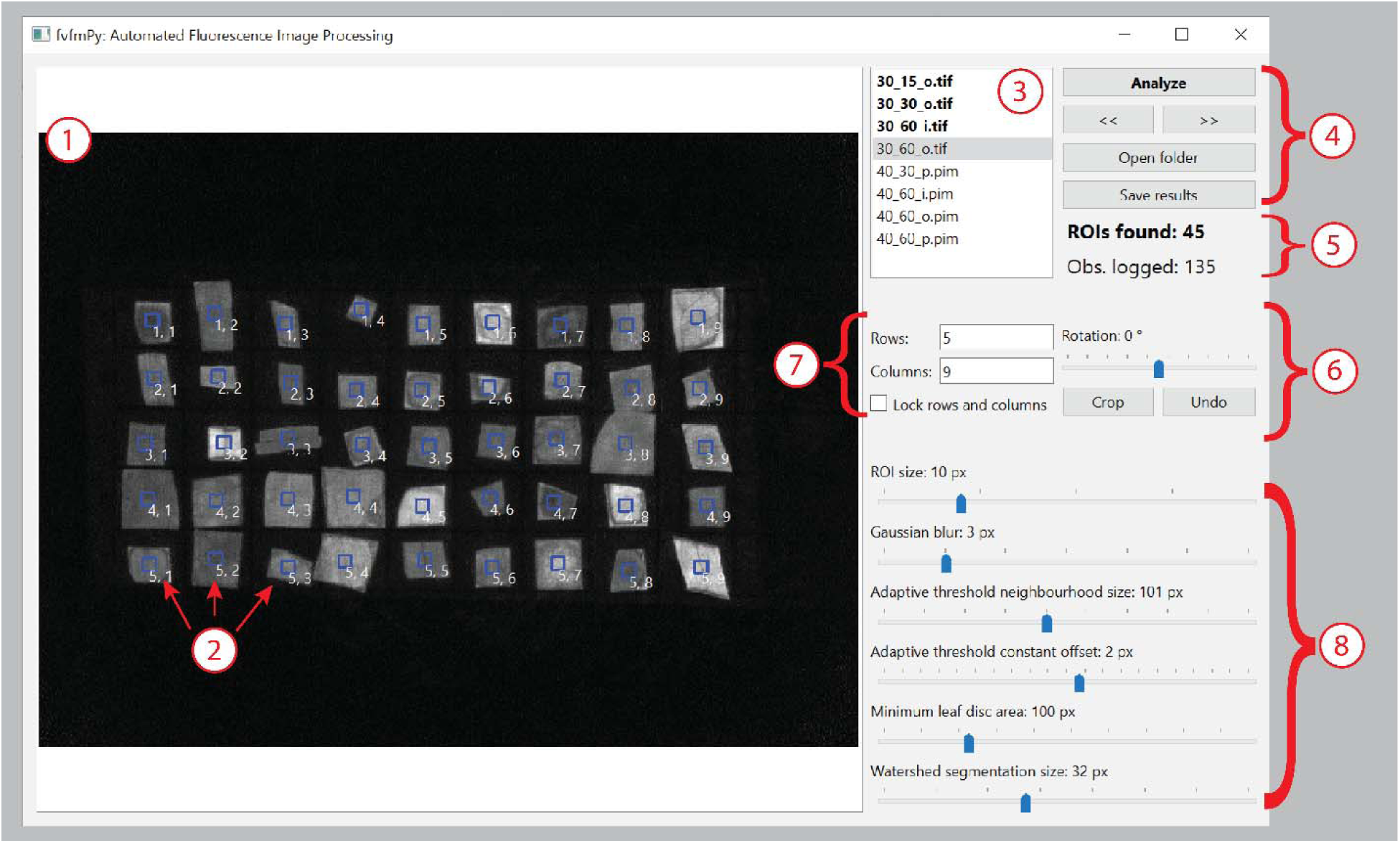
Graphical user interface. This figure depicts the graphical user interface of `fvfmPy` with a fluorescence leaf array displayed. Highlighted numbers indicate features of interest: (1) Image display. (2) Leaf discs with identified regions of interest (ROIs) overlaid in blue; row and column numbers are printed next to each ROI. (3) List of files to be analyzed in the current directory, with current file highlighted. (4) File management controls. (5) Number of ROIs identified in the current image and total number of observations recorded (total ROIs across all images analyzed so far). (6) Crop and rotate functions. (7) Number of rows and columns identified or specified by the user. (8) Sliders to adjust image segmentation parameters.

**Table 1:** Output file format.

| Column name | Description | Units or datatype |
| --- | --- | --- |
| filename | Name of processed fluorescence datafile (.pim or .tif) | String |
| row | Row number of leaf disc array | Integer |
| col | Column number of leaf disc array | Integer |
| leaf_number | Unique number of leaf disc array within file | Integer |
| centroid_x | X-coordinate of centroid of ROI | Pixels |
| centroid_y | Y-coordinate of centroid of ROI | Pixels |
| mean_Fo | Mean $F_o$ within ROI | Uncalibrated fluorescence value (unitless) |
| mean_Fm | Mean $F_m$ within ROI | Uncalibrated fluorescence value (unitless) |
| FvFm | Mean $F_v/F_m$ within ROI | Ratio (unitless) |

### Image segmentation and automated ROI selection

`fvfmPy` uses a series of algorithmic image processing steps to identify ROIs in fluorescence images of leaf disc arrays. The image processing steps employed in this software were selected to optimize the performance of the algorithm in detecting and segmenting small leaf discs in rectangular arrays. The image processing steps proceed in the following order:

1. **Gaussian blur.** A low-pass blurring filter is applied to the whole image to reduce high-frequency noise and smooth edges. The size of the blurring kernel is user-adjustable.
2. **Adaptive intensity threshold.** The algorithm next applies a Gaussian adaptive threshold to the blurred image. This works similarly to a standard brightness or colour-based threshold to separate background (i.e. cardstock) from foreground (i.e. leaf discs). However, the adaptive thresholding works on only a small “neighbourhood” of pixels at a time, rather than the entire image. This helps to prevent miscategorising leaf discs with low absolute fluorescence intensity as background. The size of the adaptive threshold neighbourhood and a constant offset value are user-adjustable.
3. **Size filtering.** The identified foreground from the previous step is then subject to a minimum size filter. Any foreground objects smaller than a threshold size are discarded. This helps to remove debris from the field of view, as well as removing small “satellite” regions of actual leaf discs (i.e., regions which appear detached from the majority of the disc due to intervening areas of low fluorescence). The minimum size threshold is user-adjustable.
4. **Watershed segmentation.** This step involves distance-transforming the foreground mask obtained from the previous steps. For each point in the foreground (i.e. pixel representing leaf material), the distance to the nearest background pixel is calculated. Local maxima in this distance-transformed space are interpreted as centres of leaf discs. This step helps to separate leaf discs which may be partially overlapping or touching and hence cannot be reliably segmented based solely on separating background from foreground. A minimum size parameter is employed in this step to prevent multiple ROIs being identified on a single leaf. The minimum size parameter is user-adjustable.
5. **Place ROIs.** The software then places a ROI at the centre of each leaf disc identified in the previous step. The size of the ROI is user-adjustable.
6. **Identify array dimensions.** Using the coordinates of the ROIs placed in the previous step, the software then attempts to determine the number of rows and columns in the image. This is accomplished by first removing any apparent rotation of the leaf disc array using singular value decomposition on the matrix of ROI centroid coordinates. Next, likely breaks between individuals rows and columns are identified by sorting gaps in horizontal and vertical coordinates of ROI centroids and identifying the largest difference in gap size; gaps between centroid coordinates lying above this threshold are interpreted as likely row or column breaks. Once the number and location of rows and columns is estimated, each ROI is assigned to its nearest row and column. The user also has the option to manually specify the number of rows and columns.

Once the automated image segmentation process is complete, the identified ROIs and their assigned row and column numbers are displayed to the user (Fig. 2). At this point, the user may accept and log the fluorescence data within the identified ROIs by pressing the “Analyze” button, or they may employ one or more optional processing steps if the initial automated image segmentation is not satisfactory.

### Optional manual steps in image processing workflow

Although the automated ROI selection algorithm is in general quite robust, it may occasionally give unsatisfactory results. This may occur, for example, due to overlapping leaf discs, excessively rotated leaf arrays, large debris, or large variation in leaf disc size. In these cases, the ROI selection can be improved by several optional steps. First, the user can rotate the image such that the rows and columns of the array are approximately horizontal and vertical, respectively, which can aid in correct identification of row and column numbers. Next, the user may crop the image to remove excess background area and any debris that may be present in the cropped area. Third, the user can adjust various thresholds and parameters employed in the automated ROI selection algorithm (see previous section). The default parameter values were selected to optimize performance based on available chlorophyll fluorescence images, but different values of these parameters may be more appropriate for other datasets. Fourth, the user can specify the number of rows and columns in the image, especially if these values were not correctly estimated by the software. Finally, the user can manually move, add, or remove ROI points in the image. We recommend using the minimum amount of manual intervention necessary to achieve satisfactory ROI placements, as this improves throughput speed and reproducibility of results.

### Fluorescence calculations from ROIs and outputs

Once the user is satisfied with the ROI placements and presses the “Analyze” button, the software calculates mean values of *F_o_* (minimum fluorescence) and *F_m_* (maximum fluorescence) on a pixel-wise basis within each ROI. Mean *F_v_/F_m_* within each ROI is then calculated as

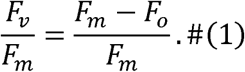

The extracted *F_o_* and *F_m_* values and computed *F_v_/F_m_* value are saved and the software advances to the next fluorescence file. Once the user has finished analysing all desired files, the results file is saved as a long-format .csv containing filename, row and column assignments, unique leaf numbers, the centroids of the identified ROIs within the image, and the extracted fluorescence metrics. The format of the results file is given in Table 1.

### Validation dataset

To ensure that the semi-automated image segmentation algorithm accurately reproduces *F_v_/F_m_* values estimated using conventional (manual) processing techniques, we performed a validation experiment. We analysed 40 chlorophyll fluorescence images comprising a total of 1799 individual *F_v_/F_m_* measurements using both manual and our new automated image processing techniques. For manual processing, one user assigned ROIs by hand using the `ImagingWIN` software associated with the fluorometer instrument (Heinz Walz GmbH, Effeltrich, Germany). For automated processing, the same user employed `fvfmPy` as described here. Manually processed *F_v_/F_m_* values were compared to automatically processed *F_v_/F_m_* values using ordinary least-squares regression in R ver. 4.4.1 (R Core Team, 2019).

### Example application: Thermal Load Sensitivity of Photosystem II

To illustrate the applicability of `fvfmPy`, we provide a complete worked example using the fluorescence data described above. We tested the Thermal Load Sensitivity (TLS) of PSII by integrating `fvfmPy` with existing analytical pipelines (Arnold *et al*., 2026, Preprint; Noble *et al*., 2026, Preprint). We selected *F_v_/F_m_* results from five Eucalypt species (de-identified; hereafter SP1-5) described above. We scaled the post and overnight *F_v_/F_m_* measurements as a proportion of the initial *F_v_/F_m_*value, such that a value of 1 indicates no functional impairment and 0 indicates total loss of function. We removed leaf samples with initial *F_v_/F_m_*< 0.5 (1% of the data), retaining a total of 895 leaf samples, each with the three corresponding *F_v_/F_m_* measurements (initial, post, and overnight).

We fit a single-stage four-parameter log-logistic (4PL) model to the *F_v_/F_m_* data that directly parameterises thermal tolerance (*CT_max_*) and thermal sensitivity (*z*) (Noble *et al*., 2026, Preprint). The 4PL model takes the form

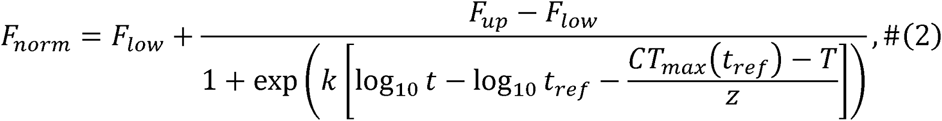

where *F_norm_*is the normalized (scaled) *F_v_/F_m_* value proportional to the initial measurement; *F_up_* is the upper asymptote; *F_low_*is the lower asymptote; *k* is a steepness metric related to the speed of decline of *F_norm_* with time; *t* is exposure duration; *t_ref_* is a reference exposure duration (fixed to 60 min in this analysis); *CT_max_*(*t_ref_*) is the critical maximum temperature at *t_ref_*; *T* is temperature; and *z* is a thermal sensitivity parameter describing the change in *T*_50_ imposed by a 10-fold change in exposure duration. Model comparisons were made with Akaike’s Information Criterion (Akaike, 1998) and were checked against convergence diagnostics and overparameterization. The best-supported 4PL model allowed *F_up_*, *CT_max_*, and *z* to vary among species, while *F_low_* and *k* were pooled among species. Plant-level repeated measures were included as a random effect on *CT_max_* only. These analyses were performed in R ver. 4.6.0 (R Core Team 2026) using the `freqTLS` R package (Nakagawa et al. 2026).

## Results

### Validation of semi-automated image segmentation method

We compared *F_v_/F_m_* values obtained using `fvfmPy` to values obtained with manual processing using ImagingWin software (Fig. 3). We found that *F_v_/F_m_* obtained using `fvfmPy` were indistinguishable from those obtained using manual processing (*r*^2^ > 0.99; slope 0.998; intercept -0.001). In addition, processing chlorophyll fluorescence images with `fvfmPy` was substantially faster than manual processing. Using `fvfmPy`, we were able to process an average of 7.54 measurements per second, or 13 seconds per 100 measurements. Manual processing, in contrast, gave an average of 0.499 measurements per second, or 200 seconds per 100 measurements. Thus, our semi-automated software pipeline reduced the time required for image analysis by approximately 94%.

**Figure 3:**
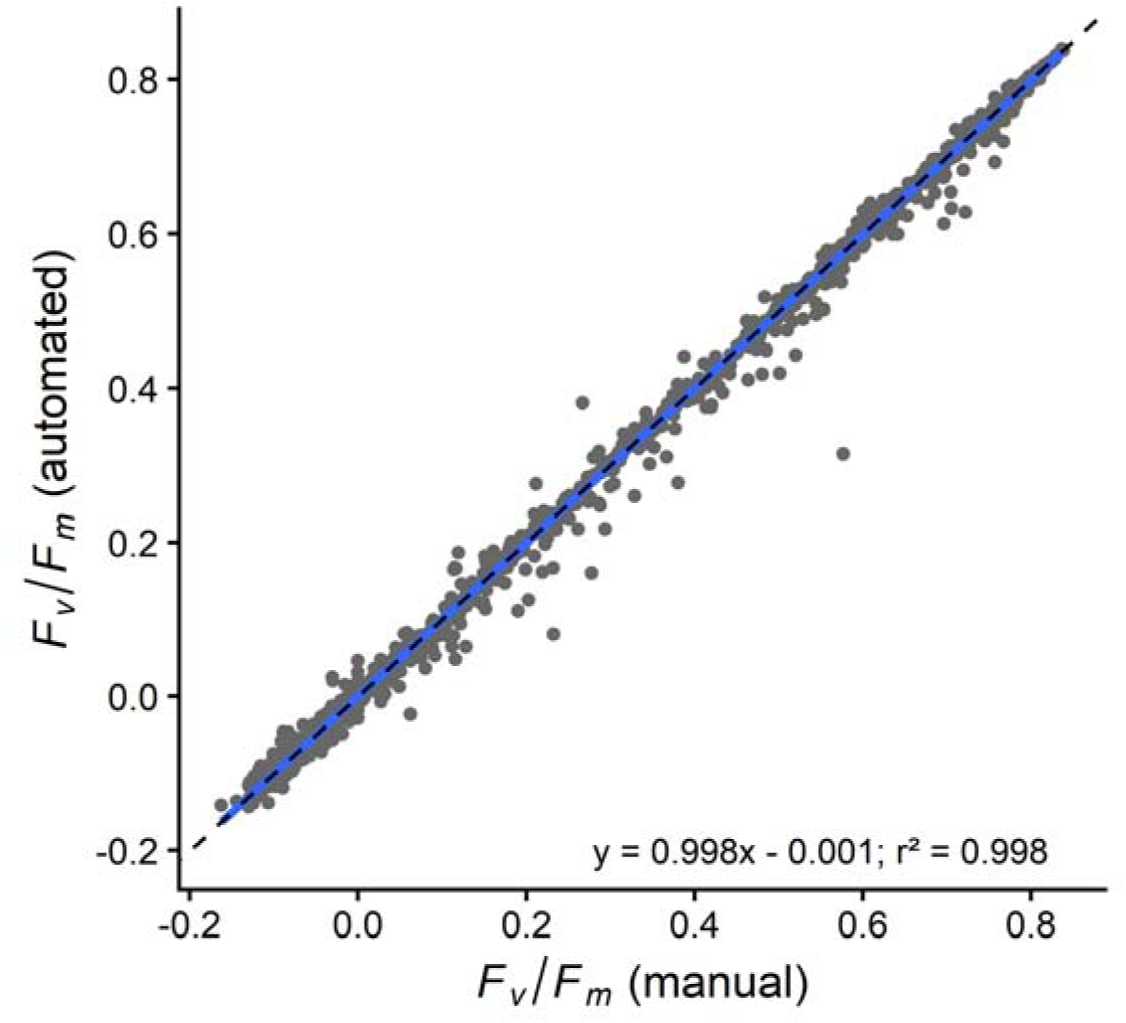
Comparison of fluorescence metrics between manual and automated processing methods. Points represent individual fluorescence measurements (*F_v_/F_m_*; *n* = 1799) which were measured using both manual processing and the automated `fvfmPy` pipeline. Blue line is ordinary least squares regression, with equation of best fit and coefficient of determination shown. Dashed black line is 1:1. Results were nearly indistinguishable between the two methods, but automated processing was completed in 94% less time than manual processing.

### Example application: Thermal Load Sensitivity of photosystem II

Our example application of the `fvfmPy` software to Thermal Load Sensitivity revealed substantial variation in critical heat tolerance thresholds and thermal sensitivity metrics (Fig. 4, Table 2). The best-supported 4PL model identified that the lower asymptote (*F_low_*) and the slope of the decline in *F_v_/F_m_* with heat dose (*k*) did not vary among species (*F_low_* = 0.031; *k* = 3.261). However, the five measured species differed significantly in heat tolerance at 1 h exposure duration (*CT_max_* ranging from 40.6 to 43.8°C), thermal sensitivity (i.e. change in *T*_50_ imposed by a 10-fold change in exposure duration; *z* ranging from 2.9 to 3.7 °C), and the upper asymptote (*F_up_* ranging from 0.71 to 0.85). See Table 2 for complete fit statistics.

**Figure 4:**
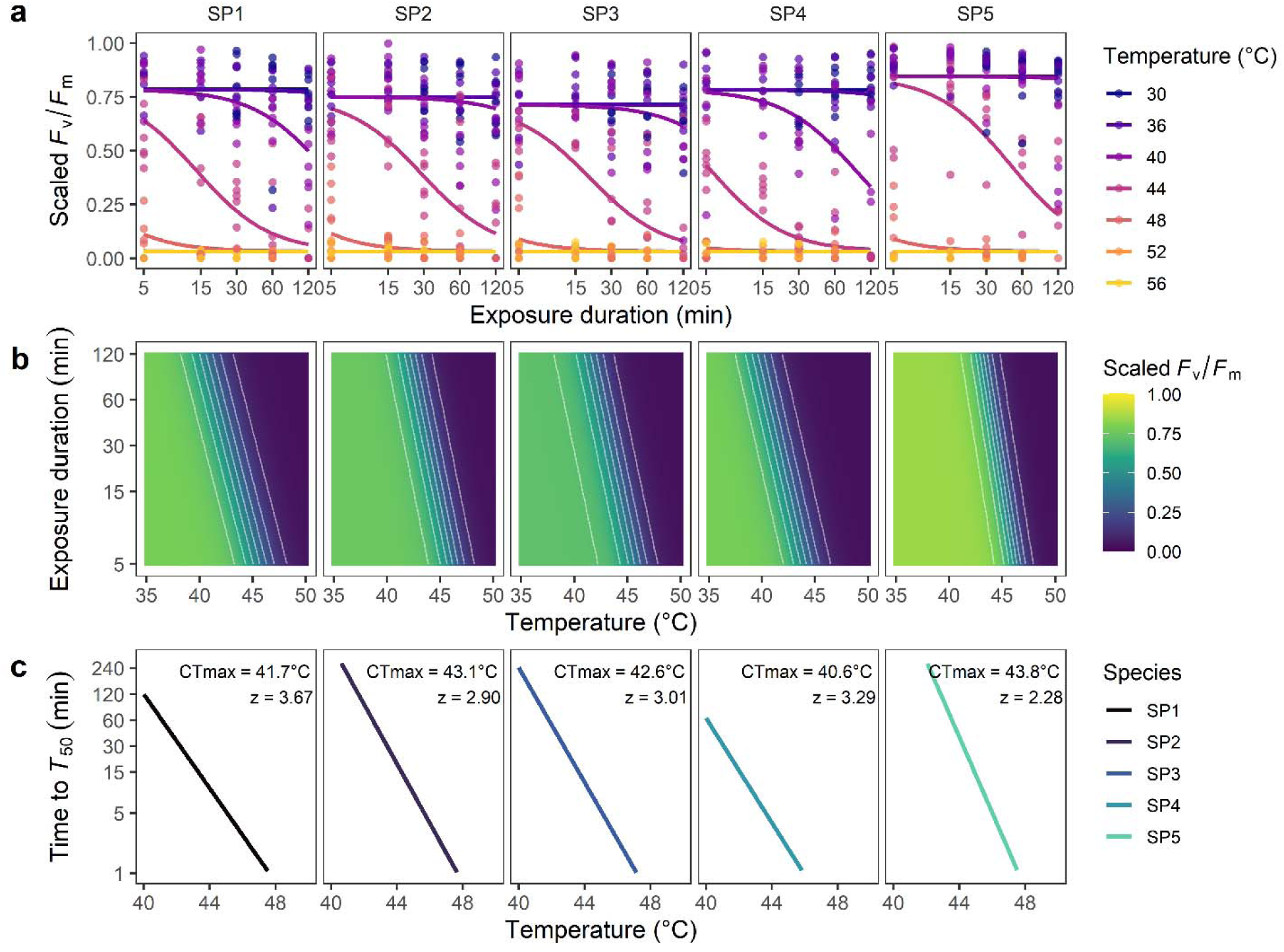
Thermal Load Sensitivity example using data extracted from the semi-automated image analysis pipeline. (a) Scaled *F_v_/F_m_*(overnight value relative to initial value) varies depending on temperature, exposure duration, and species. Coloured dose-response curves are fitted for each species and temperature combination directly from the single-stage four-parameter log-logistic model. (b) Heat dose surface of scaled *F_v_/F_m_* across exposure duration and temperature. (c) Time to reach 50% decline in *F_v_/F_m_*at a given temperature across species. Derived heat tolerance after 1 hour of exposure (*CT_max_*) and thermal sensitivity (*z*) at the *T*_50_ level are inset. Note the logarithmic scale for duration.

**Table 2:** Summary output from fitted 4PL model.

| Parameter | Species | Estimate | SE | z-value | p-value |
| --- | --- | --- | --- | --- | --- |
| $k$ | — | 3.261 | 0.273 | 11.95 | < 0.001 |
| $F_{low}$ | — | 0.031 | 0.002 | 14.06 | < 0.001 |
| $F_{up}$ | SP1 | 0.785 | 0.017 | 45.99 | < 0.001 |
|  | SP2 | 0.751 | 0.017 | 43.04 | < 0.001 |
|  | SP3 | 0.714 | 0.019 | 37.69 | < 0.001 |
|  | SP4 | 0.783 | 0.018 | 43.61 | < 0.001 |
|  | SP5 | 0.845 | 0.012 | 69.69 | < 0.001 |
| $CT_{max}$ | SP1 | 41.655 | 0.445 | 93.68 | < 0.001 |
|  | SP2 | 43.075 | 0.536 | 80.41 | < 0.001 |
|  | SP3 | 42.555 | 0.517 | 82.36 | < 0.001 |
|  | SP4 | 40.585 | 0.429 | 94.70 | < 0.001 |
|  | SP5 | 43.764 | 0.407 | 107.65 | < 0.001 |
| $z$ | SP1 | 3.671 | 0.293 | 12.54 | < 0.001 |
|  | SP2 | 2.899 | 0.508 | 5.70 | < 0.001 |
|  | SP3 | 3.011 | 0.367 | 8.21 | < 0.001 |
|  | SP4 | 3.295 | 0.274 | 12.02 | < 0.001 |
|  | SP5 | 2.277 | 0.241 | 9.46 | < 0.001 |
| plant_id | — | 0.887 | 0.137 | 6.47 | < 0.001 |

Across species measured here, scaled *F_v_/F_m_* varied depending on temperature, exposure duration, and species identity (Figure 4a). The response of scaled *F_v_/F_m_* to both temperature and exposure duration can be visualized as a heatmap, revealing species-specific differences in heat dose response and thermal sensitivity (Figure 4b). Finally, the time required to reach 50% decline in *F_v_/F_m_*at a given temperature can be visualized as a linear relationship when duration is logarithm-transformed (Figure 4c), in which the slope *z* characterizes the thermal sensitivity of the species. For example, at 44 °C, SP4 reaches 50% loss of function in < 4 min, compared to c. 10 min for SP1, and c. 38 min for SP5. Thus, a seemingly small difference in *CT_max_* and *z* translates into a large difference in tolerance capacity at a shared temperature. These results demonstrate that `fvfmPy` and `fvfmR` can integrate directly into existing data analysis pipelines for estimating PSII heat tolerance and thermal load sensitivity.

## Discussion

We developed and tested `fvfmPy` and `fvfmR`, user-friendly software pipelines written in Python and R for rapid, semi-automated processing of chlorophyll fluorescence images. `fvfmPy` drastically reduces the time required for image processing by automatically identifying leaf discs, placing regions of interest, and extracting fluorescence metrics. `fvfmPy` improves reproducibility over manual methods by 1) minimizing the need for manual user input, 2) reducing user-to-user variability, 3) reducing the potential for subjective bias in manual ROI placement, and 4) recording the locations of ROIs so that the images can be reanalyzed in an identical manner in the future. Fluorescence metrics obtained with `fvfmPy` are indistinguishable from those obtained using manual image analysis, and `fvfmPy` and `fvfmR` integrate easily with other software packages (e.g., `freqTLS` and `bayesTLS`) for seamless downstream data analysis.

### Best practices for fluorescence data collection

The `fvfmPy` software has been developed and optimized for detecting small leaf discs in a rectangular array. We have observed somewhat reduced performance when the software is used on non-rectangular arrays, leaf discs which vary greatly in size, or arrays in which leaf discs overlap or touch. Here, we offer several suggestions to keep in mind when building leaf arrays and collecting data to help improve data extraction performance.

We recommend using only rectangular grids of leaf discs. For example, the leaf array shown in Fig. 2 is a rectangular array with 5 rows and 9 columns. Any number of rows and columns may be used, provided the leaf discs are arranged into approximately straight rows and columns. If additional leaf discs are necessary (i.e. a partial row or column), we recommend adding additional “dummy leaf discs” to make a complete row or column, which may be removed from the data prior to analysis.

We recommend using clearly separated leaf discs and roughly uniform leaf disc size where possible. Although the watershed segmentation step can separate leaf discs that are touching or partially overlapping, best performance is ensured by placing leaf discs onto the array with clear gaps in between. Additionally, large differences in leaf disc size may result in suboptimal performance. For example, excessively small leaf discs may be misidentified as background, or excessively large discs may have multiple ROIs identified within them.

We also recommend ensuring that the leaf disc array is correctly aligned with the imaging fluorometer camera. Keeping the rows and columns of the leaf disc array aligned with the horizontal and vertical axes of the camera improves the ability of the algorithm to identify and assign correct rows and columns to ROIs. However, we have also provided an image rotation function to compensate if the image is excessively rotated. Following these best practices during data collection will help improve the performance of the image processing algorithm and reduce the need for manual intervention in the ROI placement process.

### Applications

`fvfmPy` and `fvfmR` have many possible applications in high-throughput chlorophyll fluorescence processing, and these tools have been developed to integrate easily into existing, commonly-used data pipelines. A key application of `fvfmPy` is in measurement of photosynthetic heat tolerance and thermal load sensitivity, as illustrated in our worked example (Fig. 4, Table 2). Measurement of PSII heat tolerance, and especially thermal load sensitivity (TLS), necessitates large quantities of data to achieve robust parameter estimates (Perez and Feeley, 2020; Arnold *et al*., 2026, Preprint). TLS, in particular, requires repeated fluorescence measurements under many temperature-by-duration treatment combinations, which may require hundreds of individual leaf disc samples to thoroughly characterize TLS for a single species.

Hence, large measurement campaigns characterizing heat tolerance or TLS for many species, genotypes, or treatments can result in extremely large datasets requiring many hours or days of manual effort to analyse and extract fluorescence metrics. Recent interest in characterizing acclimation of photosynthetic heat tolerance in response to warming or extreme heat events necessitates repeated measurements of chlorophyll fluorescence at high temporal resolution, further increasing data demands (Drake *et al*., 2026; Posch *et al*., 2026). We demonstrated that our new software has the potential to drastically reduce measurement time during the image processing step, improve reproducibility, and integrate with existing data processing pipelines into a complete, ultra-high-throughput workflow for TLS analysis. In our example, we integrated `fvfmPy` with the `freqTLS` R package pipeline (Nakagawa, 2026), however `fvfmPy` is also easily interoperable with other packages such as `bayesTLS` (Noble *et al*., 2026, Preprint).

High-throughput phenotyping is an active area of development in agronomy, where large field trials may involve screening hundreds or thousands of genotypes of crop species (Shakoor *et al*., 2017; Yang *et al*., 2020; Gill *et al*., 2022). Chlorophyll fluorescence has been widely adopted as a screening tool for identifying stress responses to a wide variety of potential abiotic or biotic stressors in the field, including salinity, cold, heat, water logging, or drought, for example (Sayed, 2003; Baker and Rosenqvist, 2004; Arief *et al*., 2023). Chlorophyll fluorescence can be a sensitive early indicator of stress, often exhibiting changes before visual cues are apparent. For this reason, repeated fluorescence measurements during crop development trials and crop monitoring is an important potential application of this new software (Chaerle and Straeten, 2000; Murchie and Lawson, 2013; Moustaka and Moustakas, 2023). As in TLS measurement campaigns, processing thousands of individual leaf samples during large field trials or crop monitoring can present a substantial bottleneck in data throughput. The image analysis pipeline developed here has potential application for improving throughput when phenotyping many crop genotypes or monitoring the health of large crop plantations. More generally, removing the image processing bottleneck may open new opportunities to design larger experiments, increase statistical power and robustness of parameter estimates, measure more genotypes, or improve representation within individuals, or within or among species (Furbank and Tester, 2011; Minervini *et al*., 2015).

Beyond increasing throughput and scalability of chlorophyll fluorescence measurements, we note that `fvfmPy` and `fvfmR` have the additional benefit of improving reproducibility of image analysis. Manual data extraction from chlorophyll fluorescence images can be influenced by subjective bias in ROI selection or variation in ROI placement among different users. By automating ROI placement, `fvfmPy` increases the reproducibility of fluorescence metrics extracted from images. Further, the ROI placements are recorded with the output data, ensuring that the images can be reanalyzed in an identical manner in the future. Reproducibility of analyses has been identified as a key issue in ecological studies, hence tools such as `fvfmPy` are needed (Powers and Hampton, 2019).

### Limitations and future directions

Chlorophyll fluorescence imaging is a powerful approach for detecting plant stress responses and photosynthetic performance, but its wider adoption in large-scale phenotyping studies depends on the availability of standardized and automated analysis pipelines (Pérez-Bueno *et al*., 2019). Our new software is an important step towards high-throughput processing of chlorophyll fluorescence imaging, improving analytical efficiency, consistency, and reproducibility. While our software has been developed to process chlorophyll fluorescence images using rectangular leaf disc arrays and a particular imaging PAM system, these image-processing software packages can provide a jumping-off point for future application. For example, additional file-handling modules could easily be added to our pipeline to improve interoperability and support for other instruments (e.g., the Photon Systems Instruments FluoroCam). Similarly, the image segmentation techniques developed and implemented here are generalizable to other kinds of data, for example RGB or multispectral information may be obtained in a similar manner using an appropriate camera. Diverse fluorescence metrics are also obtainable from other PAM measurements using this same workflow to automatically identify ROIs, including dark-light induction (Kautsky curves), rapid light curves, and nonphotochemical quenching. These examples demonstrate the potential for many extensions of our software, which is open-source and freely available for use and modification.

Our automated ROI detection approach could be further expanded and adapted to detect and extract data collected *in situ*, such as imagery obtained using UAVs (unmanned aerial vehicles or drones) or satellite imagery from remote sensing. Such platforms can capture phenotypic variation in large populations across plant development stages, generating extremely large datasets (Naito *et al*., 2017; Danzi *et al*., 2019). Image analysis and distillation of such complex datasets create even greater hurdles for data processing, and adapting automated image analysis to such situations will likely require machine learning and artificial intelligence approaches (Wang *et al*., 2026). For example, convolutional neural networks show great potential for unsupervised image segmentation and feature extraction (Xu *et al*., 2021). Applying deep learning models to plant phenotyping tasks can distil insights from highly complex and multidimensional data, but the performance of such models is greatly dependent on the quality and diversity of the dataset, and generalizability remains a concern (Singh *et al*., 2018). Field-based imagery also introduces additional challenges, as trait estimates are often sensitive to the light conditions in which the image was taken, background debris, and sensor calibration, potentially affecting the accuracy and comparability of measurements among experiments (Fahlgren *et al*., 2015; Yang *et al*., 2020; Danilevicz *et al*., 2021). Thus, high-throughput phenotyping, including chlorophyll fluorescence-based imaging, is constrained by the need for transferable analytical workflows capable of handling diverse datasets, while maintaining reproducibility and scalability across research groups and platforms (Yang *et al*., 2020).

## Conclusion

Rapid phenotyping of photosynthetic traits, including stress responses and heat tolerance, remains challenging due to the complexity of the photosynthetic apparatus and the logistical challenges associated with measurement. Enhancing measurement throughput and analytical reproducibility demands new solutions for chlorophyll fluorescence image processing. Here, we have developed a purpose-built application for analysing chlorophyll fluorescence images of leaf arrays in a semi-automated, rapid, and reproducible manner, relieving a substantial bottleneck in chlorophyll fluorescence imaging. We anticipate that advances such as this will enhance the capacity of image-based phenotyping platforms to support large-scale investigations of plant adaptation and resilience under future climate scenarios (Yang et al., 2020; Danilevicz et al., 2021; Wang et al., 2025).

## Acknowledgements

The authors acknowledge the use of generative artificial intelligence products for assistance with programming tasks (Anthropic Claude Code 2.1.25, OpenAI ChatGPT-5.6 Luna, and Google Gemini, no version number). All AI-generated code was thoroughly reviewed and tested by the authors, and the authors bear ultimate responsibility for the content of all code associated with this project.

## Author Contributions

JCG conceived the project and developed the core image analysis algorithm. JCG and PAA developed additional software features and the user interface. All authors contributed to data collection, data analysis, and writing and revising the manuscript.

## Data Availability

`fvfmPy` is published and freely available from The Python Package Index (PyPI; https://pypi.org/project/fvfmpy). Source code for `fvfmPy` is publicly available at https://github.com/garenj/fvfmPy. The R wrapper package `fvfmR` can be installed from https://github.com/garenj/fvfmR, and source code for `fvfmR` and example fluorescence images are publicly available at the same URL. `fvfmR` has also been submitted to the Comprehensive R Archive Network (CRAN; https://cran.r-project.org/) and the URL for this package will be added once approved. Data and code used to produce the data-based figures and tables in this paper are available at https://github.com/garenj/fvfmPy.figures.

## Abbreviations

*CT_max_*: Maximum critical temperature
*F_low_*: Lower fluorescence asymptote
*F_up_*: Upper fluorescence asymptote
*F_v_/F_m_*: Maximum quantum yield of PSII
*k*: Slope parameter
PSII: Photosystem II
ROI: Region of interest
TLS: Thermal load sensitivity
*T*_50_: Heat tolerance (temperature at 50% reduction of *F_v_/F_m_*)
*z*: Thermal sensitivity parameter

